# Plasma FABP4 is associated with liver disease recovery during treatment-induced clearance of chronic HCV infection

**DOI:** 10.1101/698217

**Authors:** Jean-Baptiste Gorin, David F. G. Malone, Benedikt Strunz, Tony Carlsson, Soo Aleman, Niklas K. Björkström, Karolin Falconer, Johan K. Sandberg

**Author notes:** Corresponding author: Dr. Johan K. Sandberg, CIM, Department of Medicine, Karolinska Institutet, 14152 Stockholm, Sweden.

## Abstract

Direct-acting antivirals (DAAs) have dramatically improved the management of chronic hepatitis C (CHC). In this study, we investigated the effects of hepatitis C virus clearance on markers of systemic inflammation measured in plasma samples from CHC patients before, during and after DAA therapy. We identified a plasma soluble protein profile associated with CHC. Successful DAA therapy rapidly normalised the plasma inflammatory milieu, with the notable exception of soluble (s) CD163, a marker of macrophage activation, which remained elevated after viral clearance and segregated patients with high and low levels of cirrhosis. Patients who received DAA in combination with Ribavirin maintained elevated levels of CXCL10, consistent with an immune-stimulatory role of Ribavirin. As anticipated, DAA-treated patients experienced durable improvement in liver fibrosis measurements. Interestingly, pre-treatment levels of fatty acid-binding protein 4 (FABP4) were inversely associated with reduction of APRI and FIB-4 scores during treatment. Together, these results support the notion of a rapid restoration of many aspects of the inflammatory state in CHC patients in response to DAA therapy. Furthermore, the associations with sCD163 and FABP4 warrants further investigation into the role of macrophages in residual liver disease and fibrosis resolution after viral clearance.

## Introduction

Over 70 million individuals are infected with Hepatitis C virus (HCV) worldwide. ^1^ During the last decade, direct acting antivirals (DAA) have become available for treatment of chronic hepatitis C (CHC) and have dramatically improved clinical outcomes. DAA combinations have proven to be highly efficient, allowing for sustained virologic response (SVR) rates approaching 100%,^2^ with a standard treatment course of 12 weeks that can even be shortened to 8 weeks in some instances. ^3^ These treatments have sparked hopes of eradicating HCV and the WHO has set the goal of a 90% reduction in new cases of chronic infection by 2030. ^4^ Nevertheless, even after successful elimination of HCV, risks of residual liver disease and development of hepatocellular carcinoma (HCC) remain, ^5^ It is therefore important to understand the dynamics of inflammation and fibrosis resolution during and after successful DAA treatment.

Although HCV infection localises to the liver, chronic hepatic inflammation causes systemic changes in blood cytokine and chemokine levels. ^6,7^ Numerous studies have investigated whether plasma levels of such soluble markers could predict clinical outcome of interferon (IFN)α-based therapy. ^8,9^ In the present study, we aimed to characterise how viral clearance affects plasmatic levels of cytokines and inflammatory markers in a cohort of CHC patients successfully treated with DAA. We hypothesised that some of these plasmatic proteins may be associated with the evolution of liver disease and we therefore investigated associations with clinical measures of liver status such as liver stiffness measurement (LSM), Aspartate transaminase-to-platelet ratio Index (APRI) ^10^ and Fibrosis index based on 4 factors (FIB-4) ^11^ scores.

## Results

### Plasma cytokine alterations associated with CHC and evolution during DAA therapy

To investigate the impact of HCV clearance on the immune system, 28 CHC patients were sampled before, during and after successful DAA treatment. In addition, blood samples from 20 healthy donors (HD), and 12 patients suffering from alcohol-induced cirrhosis (AC) were included as non-HCV infected comparison groups (Table 1). Plasma concentrations of 25 soluble factors, all known to be related to the immune response or inflammation, were measured in each plasma sample (Supplementary Dataset online). Out of the 25 soluble factors measured, four were significantly different (p < 0.05) between HD and CHC patients who were going to initiate IFN-free DAA therapy: CCL5, CXCL10, sCD14 and sCD163 (Fig. 1a). CCL5 was detected at lower levels (p = 0.0095), whereas CXCL10, sCD14 and sCD163 were higher in the CHC patients compared to HD (p < 0.0001, p = 0.0095 and p < 0.0001, respectively). Interestingly, there were no significant differences in CCL5 and CXCL10 levels between CHC patients and AC patients (p = 0.4764 and 0.0964) but sCD14 and sCD163 were significantly higher in CHC patients compared to AC (p = 0.0346 and 0.0489, respectively; Fig. 1a).

**Table 1:** Cohort baseline characteristics. Values are count (and percentage) for categorical variables and median [IQR] for continuous variables. ND: Not determined

|  | *HD* | *AC* | *DAA* |
| --- | --- | --- | --- |
|  | n = 20 | n = 12 | n = 28 |
| *M / F* | 50% / 50% | 92% / 8% | 50% / 50% |
| *Age (years)* | 57.00 [7.75] | 60.00 [12.25] | 60.50 [11.25] |
| <sup>2</sup><br>*BMI (kg/m )* | ND | 30.77 [4.38] | 25.94 [5.21] |
| *Diabetes* | ND | 4 (33%) | 4 (14%) |
| *Alcoholic* | ND | 12 (100%) | 11 (39%) |
| *Cirrhosis* | ND | 12 (100%) | 25 (89%) |
| *Child Pugh A / B / C* | - | 6 / 5 / 1 | 18 / 7 / 0 |
| <sup>6</sup><br>*Viral load (10 /mL)* | - | - | 1.06 [3.18] |
| *Genotype 1/2/3/4* | - | - | 19 / 2 / 6 / 1 |
| *Prior treatment* | - | - | 18 (64%) |
| *Albumin (g/L)* | ND | 33.00 [8.75] | 34.50 [8.25] |
| *Bilirubin (μmol/L)* | ND | 18.50 [24.50] | 14.00 [13.25] |
| *ALT (μkat/L)* | ND | 0.54 [0.29] | 1.27 [0.97] |
| *AST (μkat/L)* | ND | 0.82 [0.36] | 1.37 [1.04] |
| *PT (INR)* | ND | 1.20 [0.23] | 1.15 [0.22] |
| <sup>9</sup><br>*Thrombocytes (10<sup>9</sup>/L)* | ND | 140.00 [70.00] | 127.50 [83.50] |
| *LSM (kPa)* | ND | 24.35 [19.25] | 21.70 [16.25] |
| *APRI score* | ND | 0.90 [0.95] | 1.92 [1.99] |
| *FIB-4 index* | ND | 4.43 [3.25] | 6.22 [5.35] |

**Fig. 1.**
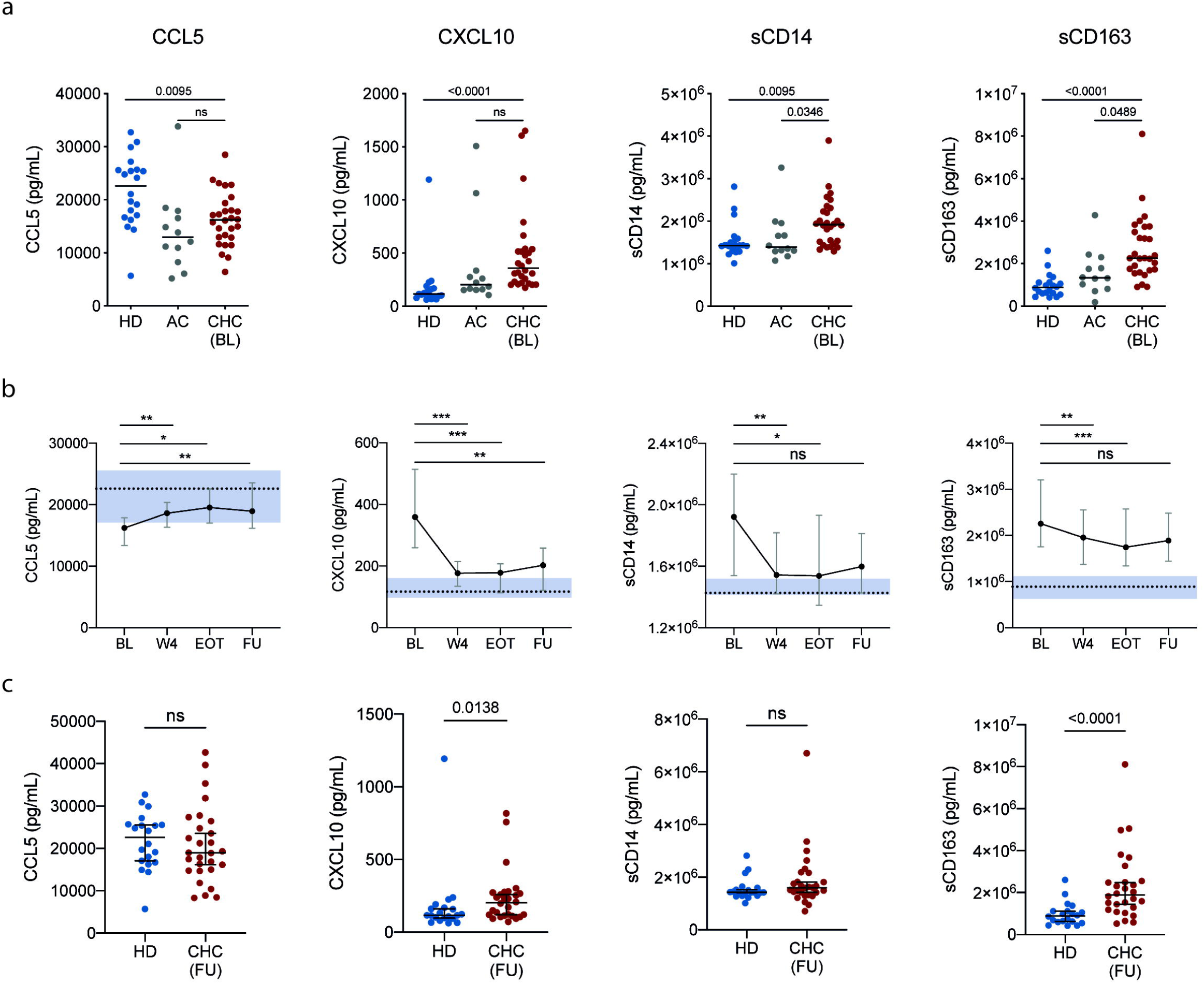
Baseline differences and evolution of plasma cytokines between healthy controls and patients. (a) Comparison between, healthy donors (HD), alcoholic cirrhotics (AC) and chronic hepatitis C patients before they initiated DAA treatment (DAA). (b) Evolution of plasma cytokine levels (median ± IQR) during DAA therapy. Dotted lines and shaded area represent the median and 95% confidence interval of the median for healthy donors. Stars represent significant changes when compared with baseline levels in CHC patients * p < 0.05; ** p < 0.01; *** p < 0.001; **** p < 0.0001. (c) Comparison between HD and CHC 6 months after the end of DAA treatment (FU).

Upon start of DAA therapy, differences with HD were quickly reduced (Fig. 1b), in contrast to the pattern observed in a previously treated cohort of CHC patients who received pegylated-IFN therapy and successfully achieved SVR (Supplementary Fig. S1 online). At follow-up (FU) after completion of DAA therapy, both CCL5 and sCD14 were restored to levels comparable to HD, whereas CXCL10 and sCD163, although significantly decreased compared to BL, remained significantly elevated in comparison to HD (p = 0,0138 and p < 0.0001, respectively; Fig. 1c).

### Effect of Ribavirin on plasma cytokine profiles

In our cohort, half of the patients who underwent DAA therapy received RBV as part of their treatment regimen (Supplementary Fig. 2 online), we therefore stratified CHC patients according to the use of RBV to assess possible effects on plasma cytokine levels (Fig. 2). There were no other significant differences before treatment between these two subgroups regarding age, BMI, liver health indicators (ALT, AST, albumin, bilirubin, PT-INR, thrombocytes, LSM) or any of the measured soluble markers. The patients that received RBV-free therapy quickly normalised all soluble factors in their plasma except for sCD163, which remained elevated throughout treatment. In contrast, patients that received RBV in combination with DAA maintained elevated levels of both CXCL10 and sCD163 (Fig. 2a and 2b). Six month after the end of therapy, sCD163 remained significantly elevated in both subgroups (p = 0.0158 and p = 0.0004), whereas CXCL10 was significantly elevated only in the group that received RBV (p = 0.0012, Fig. 2c).

**Fig. 2.**
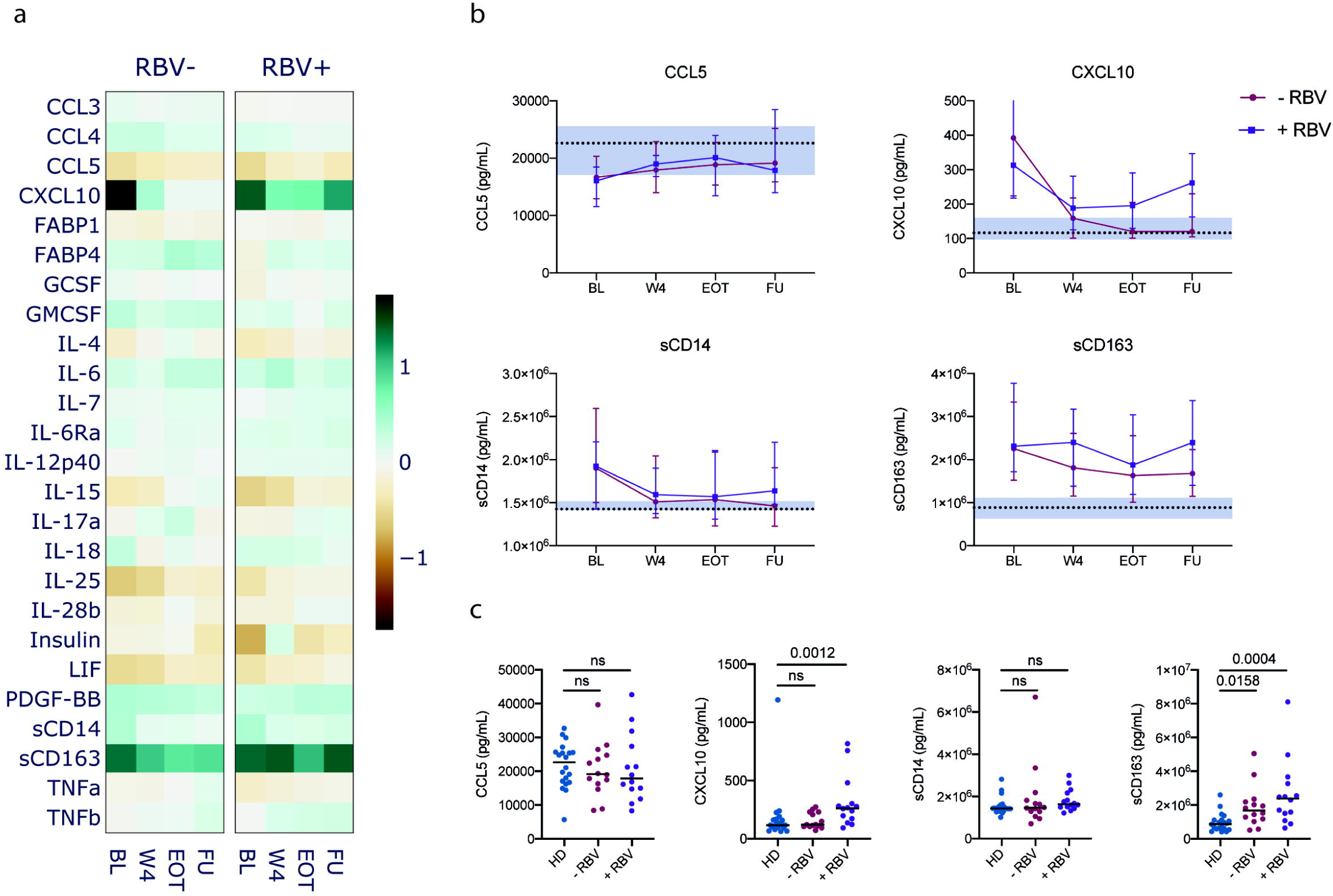
Effect of Ribavirin on cytokine levels.(a) Heatmap of median fold change of soluble marker levels in plasma compared to healthy donors for all measured cytokines in CHC patients who received a DAA regimen with (+ RBV) or without (- RBV) Ribavirin. (b) Evolution of markers concentration (median ± IQR) in CHC patients over the course of treatment. Dotted lines and shaded area represent the median and 95% confidence interval of the median for healthy donors. (c) Comparison between healthy donors (HD) and CHC patients who received a DAA regimen with (+ RBV) or without (- RBV) Ribavirin. BL: baseline; W4: week 4; EOT: end of treatment; FU: follow-up.

### Levels of sCD163 distinguish patients with different degrees of liver cirrhosis throughout DAA therapy

To assess whether the stage of liver disease was associated with the plasmatic protein profile, we stratified cirrhotic patients in the DAA group according to Child Pugh class. There were 18 patients classified as Child Pugh A and 7 patients classified as Child Pugh B. Although differences with HD seemed more pronounced in the Child B group (Fig. 3a), those differences were also reduced upon start of therapy, with the exception of sCD163, which remained elevated and significantly different between the 2 subgroups throughout treatment (Fig. 3b). Even 6 months after successful therapy, sCD163 remained elevated in Child Pugh A patients (p = 0.0103) and even higher in Child Pugh B patients (p < 0.0001, Fig. 3c). We also investigated possible associations between levels of sCD163 at the FU time point, and various indicators of liver health and found significant correlations with AST levels (ρ = 0.44, p = 0.0198), Albumin (ρ = −.48, p = 0.0101) and prothrombin time (PT-INR, ρ = 0.41, p = 0.0301, Fig. 3d). Thus elevated sCD163 seems associated with liver disease and remains elevated in patients with advanced cirrhosis despite successful clearance of HCV with DAA therapy.

**Fig. 3.**
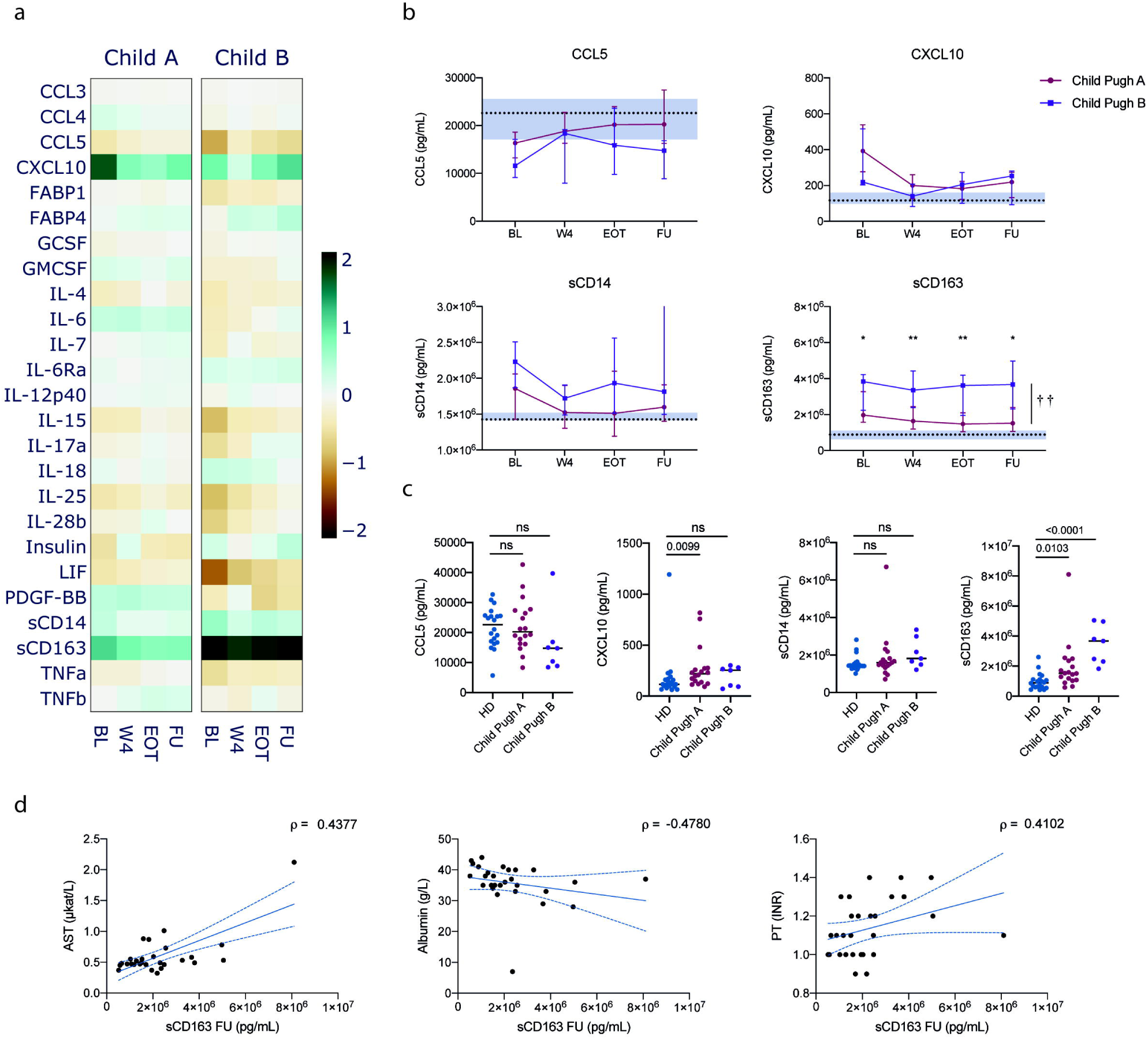
Influence of cirrhosis grade on cytokine levels over the course of therapy. (a) Heatmap of median fold change of soluble marker levels in plasma compared to healthy donors for all measured cytokines in CHC patients with cirrhosis classified as Child Pugh A (Child A) or Child Pugh B (Child B). (b) Evolution of markers concentration (median ± IQR) in CHC patients over the course of treatment. Dotted lines and shaded area represent the median and 95% confidence interval of the median for healthy donors. Crosses and stars indicate significant difference between Child Pugh A and B groups overall and for each timepoint, respectively. †† p < 0.01; * p < 0.05; ** p < 0.01. (c) Comparison between healthy donors (HD) and CHC patients with cirrhosis. BL: baseline; W4: week 4; EOT: end of treatment; FU: follow-up. (d) Spearman correlations between follow-up plasma concentration of sCD163 and different indicators of liver health. Dotted blue lines indicate 95% confidence intervals.

### IFN-free DAA combinations improve fibrosis indicators

Liver stiffness measurements were available from time points before and after therapy in 18 of the 28 DAA-treated patients (two non-cirrhotic, 14 Child Pugh A and two Child Pugh B). Mixed effects models of paired LSMs demonstrated a significant decrease in stiffness values (p < 0.0001), between baseline and follow-up measurements in the DAA group. The decrease was significant in the Child Pugh A subgroup of patients (p = 0.0003), and there was a similar trend in Child Pugh B patients (Fig. 4a). This pattern indicated decreased liver fibrosis and inflammation. Since DAA therapy efficiently eliminates the virus and quickly normalises most aspects of the cytokine milieu in plasma, it is likely that this treatment reduces liver inflammation and a decrease in LSM may thus be insufficient to conclude that there is a decrease in fibrosis. We therefore calculated APRI and FIB-4 scores for all treated patients. Both APRI and FIB-4 decreased significantly by EOT (p < 0.0068 and p = 0.0038, respectively), and this was maintained at follow-up (p = 0.0061 and p = 0.0010, respectively) (Fig. 4b and 4c). This pattern was present for Child Pugh A patient subgroup and a similar trend was observed in Child Pugh B patients (Fig. 4b-c). This is consistent with the notion that DAA therapy reduces liver fibrosis, including in patients with advanced cirrhosis. In support of these results, six out of seven patients classified as Child Pugh B before the start of therapy were re-classified as Child Pugh A at follow-up, further indicating an improvement of liver status.

**Fig. 4.**
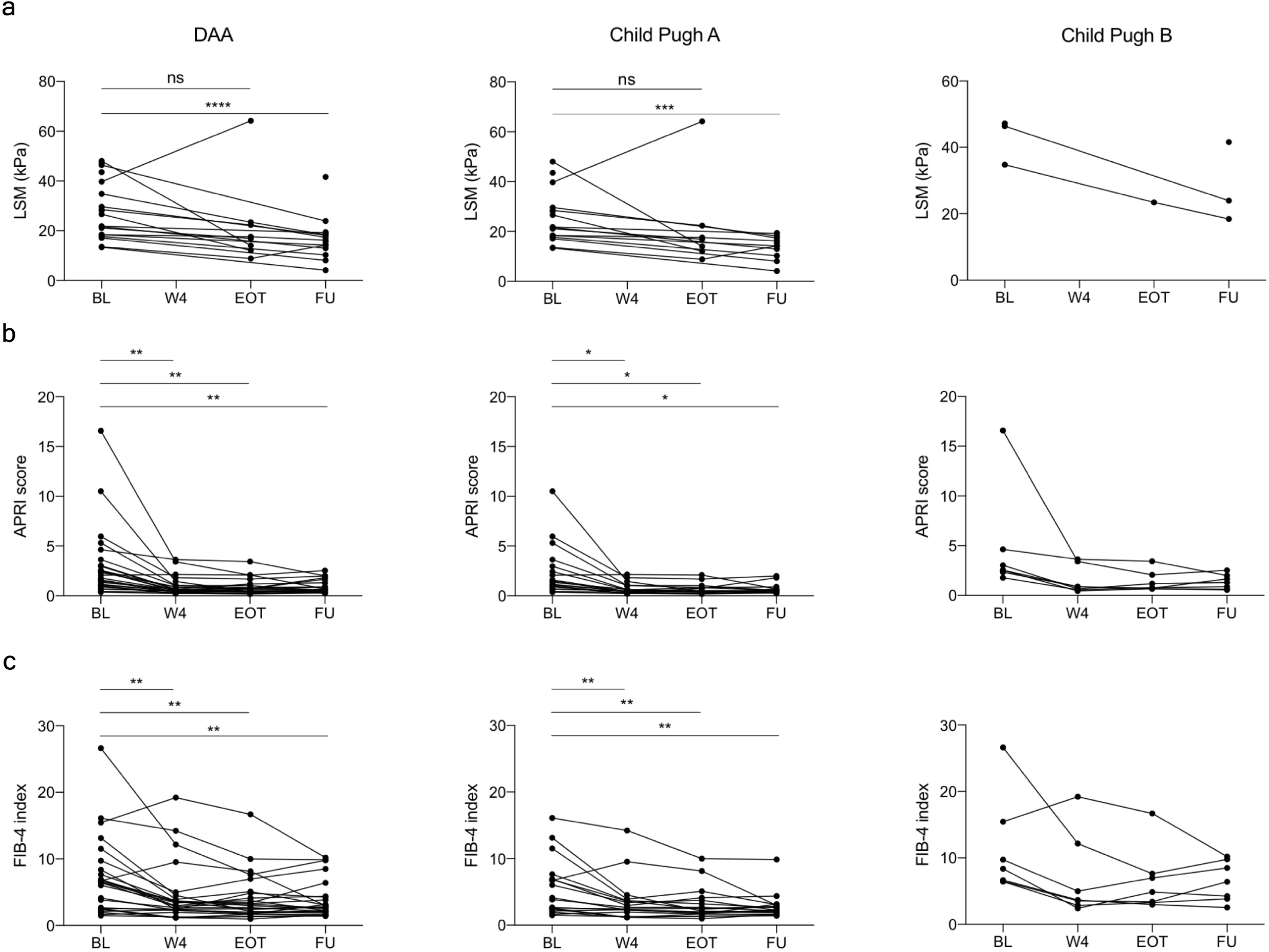
Improvement in fibrosis indicators values over the course of DAA therapy. Evolution of LSM (a), APRI score (b) and FIB-4 index (c) during the course of DAA therapy. Stars indicate statistical difference between time-points computed with repeated measurement ANOVA using mixed effect models. * p < 0.05; ** p < 0.01; *** p < 0.001; **** p < 0.0001.

### Baseline levels of soluble FABP4 are associated with improvement of fibrosis indicators in response to DAA therapy

To investigate whether soluble plasma markers might be associated with fibrosis reduction, we performed Spearman correlation analyses between baseline levels of the 25 cytokines and the reduction in LSM, APRI and FIB-4 scores observed at FU (Fig. 5). IL-18 levels at baseline correlated positively with LSM reduction (ρ = 0.54, p = 0.028), and FABP4 levels at baseline correlated negatively with reductions in both APRI (ρ = −0.53, p = 0.0035) and FIB-4 (ρ = −0.65, p = 0.0002). To investigate whether those correlations may be driven primarily by single components of the APRI and FIB-4 scores, we analysed correlations between FABP4 and platelet count increase and decrease in liver enzymes. Although FABP4 correlated significantly with platelet count increase (ρ = −0.48, p = 0.0097) and AST decrease (ρ = −0.45, p=0.0153; Supplementary Fig. S3 online), those correlations were weaker than the correlation with both composite scores, suggesting that FABP4 levels are associated to the reduction in liver disease.

**Fig. 5.**
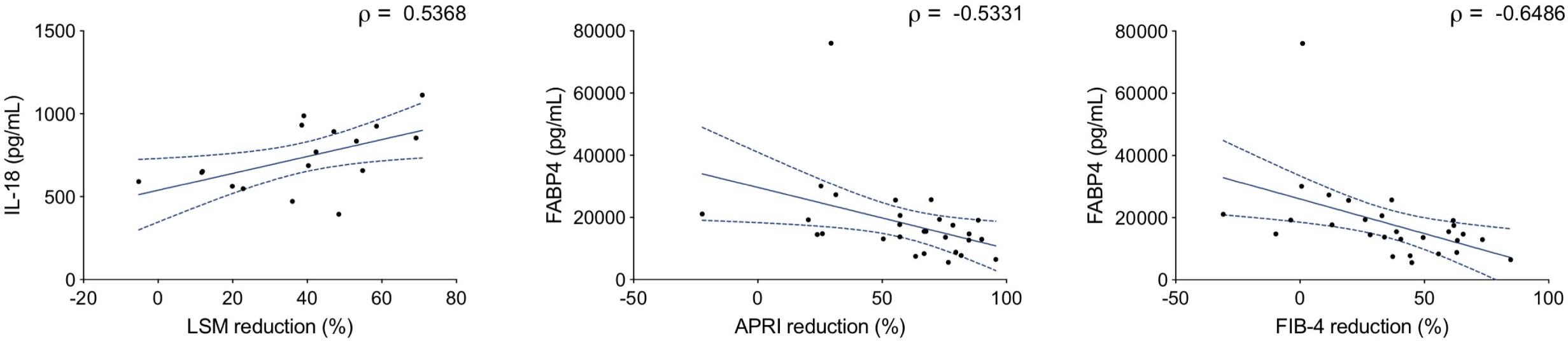
Correlations between baseline cytokine levels and decrease in fibrosis indicators. Spearman correlations between baseline plasma concentration and the improvement in fibrosis indicator values over the course of DAA treatment were computed for all cytokines. Correlations that were statistically significant (p < 0.05) are shown here. Dotted blue lines indicate 95% confidence intervals.

## Discussion

Residual liver disease and risk of developing HCC are concerns in patients that have cleared their HCV infection, and it is therefore important to understand the dynamics of systemic and hepatic inflammation during and after successful DAA treatment. Here, we identify differences in the plasmatic cytokine milieu of HD and CHC patients and show that DAA therapy rapidly reduces those differences with the exception of sCD163, which distinguishes patients with different levels of cirrhosis. We also observe that patients who receive RBV seem to maintain elevated plasma levels of CXCL10, consistent with an immune-stimulatory role of RBV. Finally, we find a significant association between baseline levels of FABP4 and subsequent reduction in APRI and FIB-4 measurements during DAA treatment, suggesting a link between FABP4 plasma levels and liver disease reduction.

Our data set identified clear differences in plasma inflammatory markers between CHC patients and HD. CCL5 was reduced in CHC patients whereas CXCL10, sCD14 and sCD163 were elevated. CXCL10 has been previously associated with viral load and it is known that HCV triggers its production through NF-K B activation. ^12^ Soluble CD14 is a marker of monocyte activation, ^13^ sCD163 is a marker of macrophage activation, ^14^ and all these three markers have been associated with liver fibrosis in untreated CHC. ^15,16,17^ However, elevated sCD14 and sCD163 are not specific to HCV infection and have been previously observed in AC. ^18-20^ Interestingly, in our cohort, although both sCD14 and sCD163 were elevated in AC, they were significantly higher in CHC patients, despite a lower proportion of Child Pugh B and C patients. Thus, the higher plasma levels of sCD14 and sCD163 in CHC compared to AC might reflect a combination of anti-viral immune activation and liver disease.

In contrast to IFN-containing regimens, which induced broad changes in the cytokine milieu, DAA therapy quickly reduced the differences observed with HD. This pattern likely reflects the very different mechanisms of action of the two treatments. DAA drugs specifically interfere with viral replication, whereas IFN is immune activating and modulates the expression of hundreds of IFN-regulated genes, which themselves can influence cytokine secretion via a complex network of interactions. ^21^ These findings are in accordance with recent work by Burchill et al. that showed a rapid decrease in the expression of genes associated with inflammation in PBMCs of HCV infected patients undergoing DAA therapy. ^22^ Certain features of T-cell and NK cell phenotype and function are also restored following DAA therapy, ^23-26^ supporting the idea of a general immune system normalisation after HCV clearance. However, recent studies also indicate that other immune cell traits, such as the appearance of regulatory T cells, ^27^ NK cell receptor repertoire diversity, ^28^ and the state of the MAIT cell compartment, ^29^ are still altered after DAA therapy. Thus, the impact on the cellular immune system might in part persist after successful viral clearance. Hengst et al. observed a partial but incomplete restoration of the cytokine milieu during DAA therapy and, notably, CXCL10 remained elevated throughout treatment. ^30^ Interestingly, all patients in that cohort received RBV in combination with DAA, and our observations indicate that patients who receive RBV maintain higher levels of CXCL10. Studies have also shown that RBV had immunomodulatory effects^31,32^ and can induce ISGs in vitro. ^33,34^ In our study, only sCD163 remained significantly elevated in patients that received DAA regimen without RBV.

Levels of sCD163 were significantly decreased by DAA treatment, as has been described by others ^35,36^. However, levels remained significantly elevated in comparison to those seen in HD. This is in accordance with work by Mascia et al. showing sustained high levels of sCD163 in DAA-treated CHC patients 12 weeks after EOT. ^37^ Additionally, we observed significant correlations between levels of sCD163 at FU and multiple indicators of liver health (AST, Albumin and PT-INR). Taken together, these findings by us and others suggest that residual macrophage activation could play a role in residual liver disease despite a general dampening of inflammation (Fig 6). Of note, the last timepoint evaluated in our study was 6 months after EOT, thus further studies would be required to assess whether elevated sCD163 reflects an irreversible consequence of CHC or a slower recovery of certain immune processes.

**Fig. 6.**
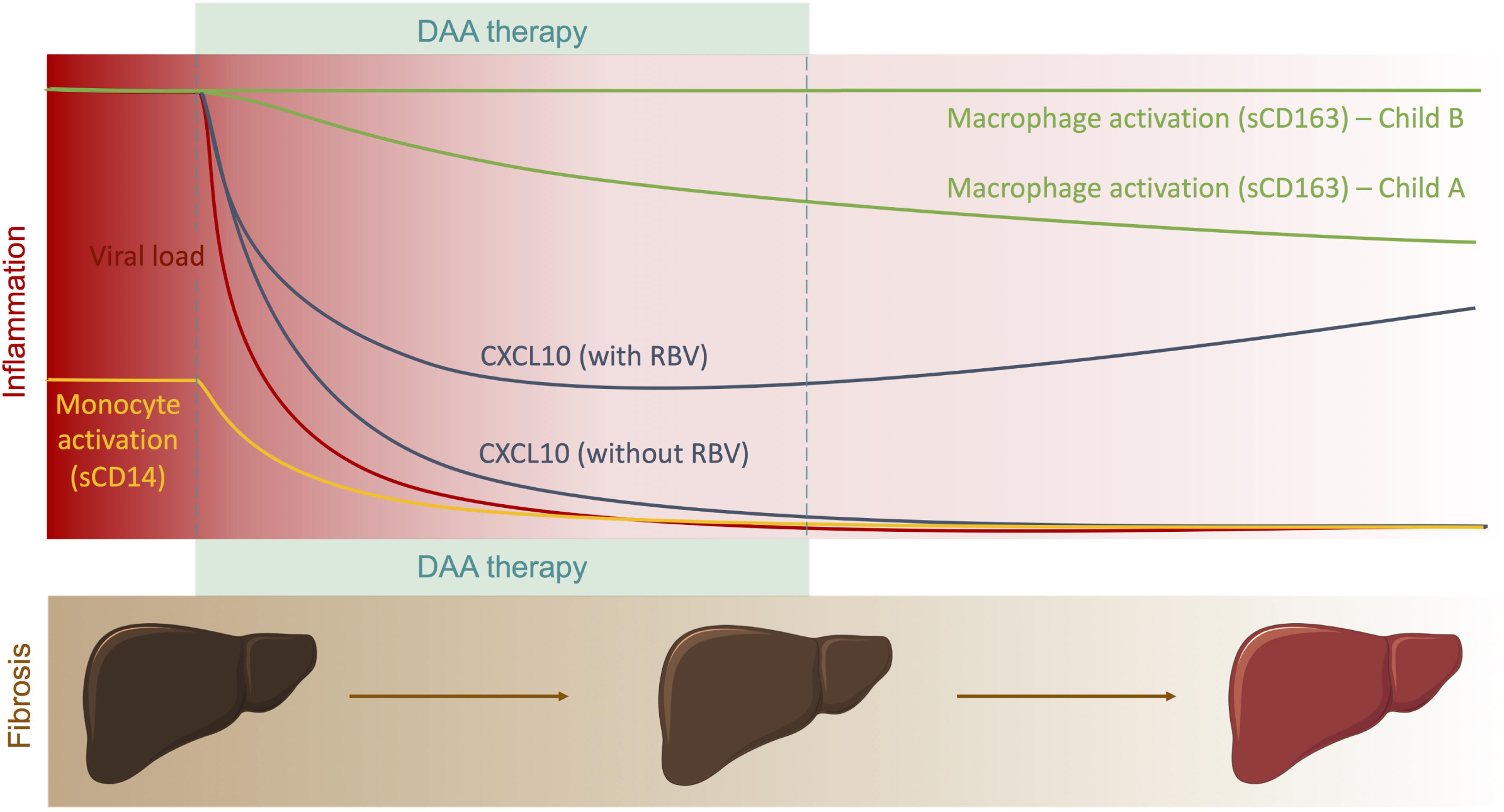
Longitudinal evolution of circulating soluble marker levels and liver fibrosis following elimination of HCV in patients undergoing DAA therapy. Liver diagrams were adapted from Servier medical art (https://smart.servier.com/) under a Creative Commons Attribution 3.0 Unported Licence (https://creativecommons.org/licenses/by/3.0/legalcode).

It is also important to understand the response of patients with more severe cirrhosis (Child Pugh B) to DAA combinations in comparison to patients with milder liver damage (Child Pugh A). Our data indicate that IFN-free DAA therapy normalises the cytokine profile, with the exception of sCD163, in both groups, and thus suggest that advanced liver damage is not a barrier to a normalised plasma cytokine milieu. Interestingly, six out of the seven Child Pugh B patients from our cohort were re-classified as Child Pugh A at the EOT, in line with improved liver function. We also observed a clear improvement in LSMs after treatment, as has been observed by others. ^38-40^ However, this measurement does not allow clear discrimination between reduction in liver inflammation, improvement of fibrosis, or a combination of the two. It has also been argued that the predictive power of liver elasticity measurements for fibrosis is reduced following therapeutic eradication of HCV. ^41^ To complement our observation, we therefore investigated the evolution of APRI and FIB-4 scores during therapy and showed a clear reduction in both of these indicators of liver fibrosis. Nevertheless, further studies comprising liver biopsies will be required to determine the extent of liver regeneration after successful DAA therapy.

We also investigated associations between plasmatic proteins and improvement of liver fibrosis and found that baseline levels of FABP4 inversely correlated with the improvement in both APRI and FIB-4 observed after treatment. FABP4 is expressed in adipocytes and macrophages, and high serum concentrations of FABP4 are associated with inflammation and risk for metabolic and vascular diseases. ^42^ Recently, increased FABP4 levels in the blood has also been shown to correlate with poor prognosis in cirrhosis. ^43^ In the same study, it was shown that FABP4 gene expression was increased in cirrhotic livers and that liver macrophages seemed to be responsible for that increase. Interestingly, sCD163 is also released by macrophages and hepatic macrophages are known to play a critical role in liver inflammation, fibrosis and resolution of inflammation. ^44^ Investigating the mechanisms leading to sCD163 and FABP4 release during CHC may thus help better understand the mechanisms of fibrosis resolution during DAA therapy. Of note, CHC cohorts are often very heterogeneous groups and further research is needed to fully understand the influence of multiple factors (such as HCV genotype, fatty liver content, obesity, diabetes, etc.) on residual inflammation and liver disease.

In conclusion, these results support the notion of a rapid restoration of the inflammatory state in CHC in response to DAA therapy and warrant further investigations of the role of macrophage activation in liver disease reduction after clearance of HCV infection.

## Methods

### Patients

To investigate the impact of HCV clearance on the immune system, CHC patients above 18 years of age scheduled to receive DAA therapy were sampled before, during and after treatment at the Department of Infectious Diseases and the Department of Gastroenterology and Hepatology at Karolinska University Hospital. Blood samples were collected from 28 CHC patients who received DAA therapy and successfully achieved SVR, and 13 CHC patients who received pegylated-IFN therapy and successfully achieved SVR. Exclusion criteria included concurrent Hepatitis B virus or HIV co-infections. In addition, blood samples from 20 HD, and 12 patients suffering from AC were included as non-HCV infected comparison groups. Of note, a proportion (64%) of patients in the DAA treated group previously underwent IFNα-based therapy without achieving SVR. The HD group was matched in gender and age to the DAA group. The study was conducted in accordance with the declaration of Helsinki, approved by the regional ethics committee (Stockholm, approval number 2012/63-31/1) and all participants gave informed consent.

### Liver Stiffness Measurement

Liver stiffness measurement was performed using transient elastography (FibroScan®). Paired LSMs before and after therapy were obtained for 18 of the 28 DAA-treated patients (2 non-cirrhotic, 14 Child Pugh A and 2 Child Pugh B liver cirrhosis patients). Measurements with an IQR > 30 or a success rate < 50% were excluded from the analysis.

### Sample collection

Venous blood was collected in heparin-coated tubes and spun 10 min at 680 g at room temperature to separate plasma, which was stored at −80°C immediately after separation. CHC patients receiving DAA therapy had blood collected at four time-points: before initiation of therapy, four weeks after initiation of therapy (W4), at the end of therapy (EOT, either 12 or 24 weeks after initiation of therapy) and at a follow-up time point collected approximatively six months after the EOT. For the IFN-treated group, 13 baseline samples, 11 W12 and 6 EOT samples (either 24 or 48 weeks after initiation of therapy) were included in the analysis.

### Luminex assay and ELISA

Plasma samples were analysed using a custom 24-plex magnetic Luminex assay (R&D systems). All samples were diluted 1:2 and assays were performed according to manufacturer’s protocol. Samples were acquired using the Bio-Plex Magpix multiplex reader (Bio-Rad) and analysed using the Bio-Plex Manager software. Soluble CD14 (sCD14) concentration in plasma samples was measured using the human CD14 Quantikine ELISA kit from R&D systems (DC140) according to manufacturer’s protocol.

### Statistical analysis

Statistical analyses were performed using Prism 8 (GraphPad). Mann-Whitney tests were used for comparisons between 2 groups and Kruskal-Wallis test with post-hoc Dunn’s correction were used for comparing more than 2 groups. Repeated measure one-way analysis of variance (ANOVA) using mixed effect models and Holm-Sidak multiple testing correction were used to analyse changes over time. Association between variables were analysed using Spearman’s rank correlation.

## Acknowledgements

We thank Britt-Mare Löfberg and Pia Loqvist for their help with collecting blood samples. We also thank the Swedish Research Council (2016-03052) and the Swedish Cancer Society (CAN 2017/777) for their financial support.

## Author Contributions

JBG collected samples, performed experiments, analysed data and co-wrote the manuscript. DFGM participated in study design, collected samples and performed experiments. BS collected samples. TC, SA and KF participated in patient recruitment. NKB, SA and KF participated in study design and data analysis. JKS coordinated the study, participated in study design, data analysis, and co-wrote the manuscript. All authors read and approved the final manuscript.

## Data Availability

All data generated or analysed during this study are included in this published article (and its Supplementary Information files).

