## Supplementary information for "Plasma FABP4 is associated with liver disease recovery during treatment-induced clearance of chronic HCV infection"

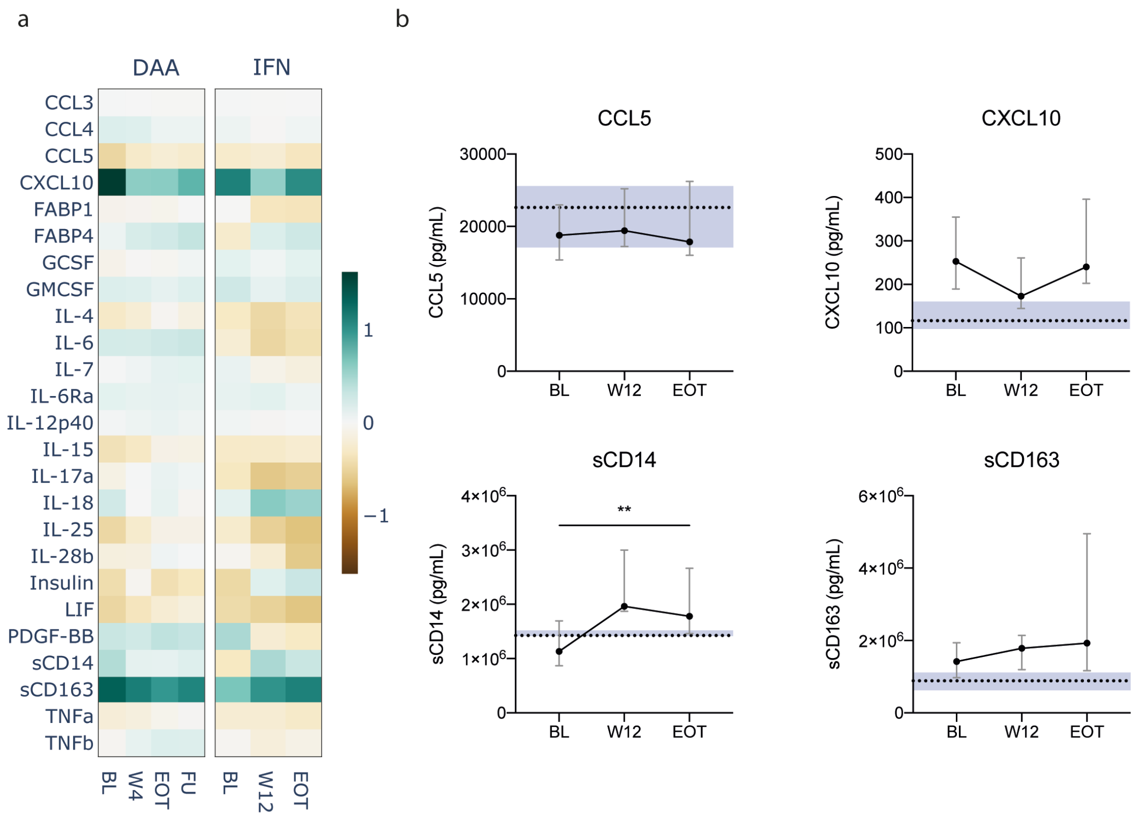


**Supplementary Fig. S1:** Comparisons between DAA and IFN therapy

(a) Heatmap of median fold change of soluble marker levels in plasma compared to healthy donors for all measured cytokines in CHC patients with cirrhosis treated with DAA or IFN regimen. (b) Evolution of markers concentration (median ± IQR) in CHC patients over the course of pegylated-IFN treatment. Dotted lines and shaded area represent the median and 95% confidence interval of the median for healthy donors. BL: baseline; W12: week 12; EOT: end of treatment.

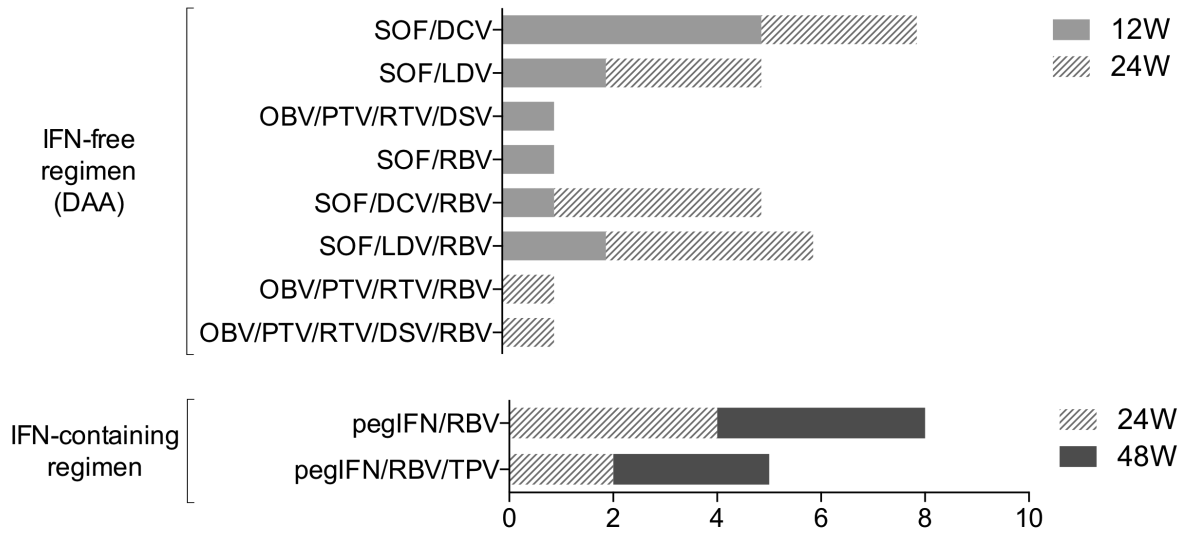


**Supplementary Fig. S2:** Treatment regimens used in the study

Histogram of number of patients assigned to each treatment regimen used in the study and the number of weeks each treatment was prescribed. DCV: Daclatasvir, DSV: Dasabuvir, LDV: Ledipasvir, OBV: Ombitasvir, PTV: Paritaprevir, RBV: Ribavirin, RTV: Ritonavir, SOF: Sofosbuvir, TPV: Telaprevir.

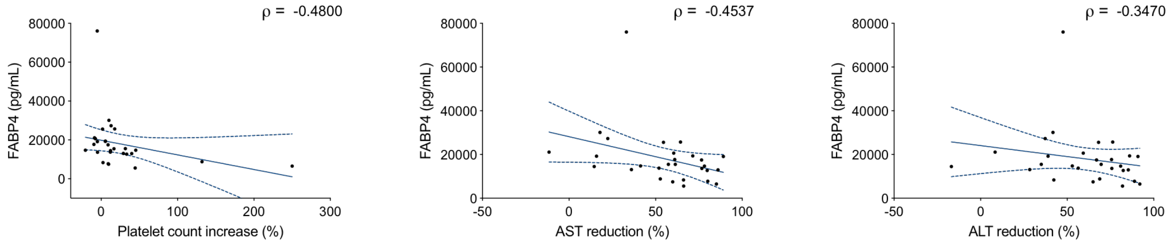


**Supplementary Fig. S3:** Correlations between baseline FABP4 and components of APRI and FIB-4 scores

Spearman correlations between baseline plasma concentration of FABP4 and the change in platelet count, AST levels and ALT levels were computed. Dotted blue lines indicate 95% confidence intervals.
